# Ecology drives the evolution of diverse siderophore-production strategies in the opportunistic human pathogen *Pseudomonas aeruginosa*

**DOI:** 10.1101/2021.01.27.427959

**Authors:** Alexandre R.T. Figueiredo, Andreas Wagner, Rolf Kümmerli

## Abstract

Bacteria often cooperate by secreting molecules that can be shared as public goods between cells. Because the production of public goods is subject to cheating by mutants that exploit the good without contributing to it, there has been great interest in elucidating the evolutionary forces that maintain cooperation. However, little is known on how bacterial cooperation evolves under conditions where cheating is unlikely of importance. Here we use experimental evolution to follow changes in the production of a model public good, the iron-scavenging siderophore pyoverdine, of the bacterium *Pseudomonas aeruginosa*. After 1200 generations of evolution in nine different environments, we observed that cheaters only reached high frequency in liquid medium with low iron availability. Conversely, when adding iron to reduce the cost of producing pyoverdine, we observed selection for pyoverdine hyper-producers. Similarly, hyper-producers also spread in populations evolved in highly viscous media, where relatedness between interacting individuals is higher. Whole-genome sequencing of evolved clones revealed that hyper-production is associated with mutations/deletions in genes encoding quorum-sensing communication systems, while cheater clones had mutations in the iron-starvation sigma factor or in pyoverdine biosynthesis genes. Our findings demonstrate that bacterial social traits can evolve rapidly in divergent directions, with particularly strong selection for increased levels of cooperation occurring in environments where individual dispersal is reduced, as predicted by social evolution theory. Moreover, we establish a regulatory link between pyoverdine production and quorum-sensing, showing that increased cooperation at one trait (pyoverdine) can be associated with the loss (quorum-sensing) of another social trait.

## Introduction

Long gone are the times when microbes were seen as solitary life forms. Over the last three decades, a wealth of research has uncovered that microbial communities are shaped by complex networks of competitive and cooperative interactions (West et al. 2007; Little et al. 2008; Mitri & Foster, 2013; Ghoul & Mitri, 2016; Granato, et al. 2019; Figueiredo & Kramer, 2020). Competitive interactions may involve the secretion of toxins against competitors, hunting or competitive exclusion (Hibbing et al. 2010; Pérez et al. 2016; Granato et al. 2019). Examples of cooperative behaviours include mutualistic cross-feeding, communication via signalling molecules, and sharing the benefits of secreted molecules such as proteases, siderophores and biofilm components (Wilder et al. 2011; D’Souza et al. 2018; Dragoš et al. 2018; Kramer et al. 2020; Robinson et al. 2020). Microbial cooperation often underlies important biological processes, including the establishment of infections (Ackermann et al. 2008; Granato et al. 2018), nutrient fixation in the rhizosphere (Denison et al. 2003) and mutualistic interactions with hosts (Verma and Miyashiro 2013).

Microbial cooperation has attracted the attention of evolutionary biologists not only because of its variety in form and function, but also because it incurs a cost for the actor while benefiting other individuals (West et al. 2007). Cooperation could thus select for “cheating” variants that do not cooperate themselves, but benefit from cooperative acts performed by others (Ghoul et al. 2014). How then can cooperative traits be maintained on evolutionary timescales? This question has spurred an enormous body of research focussing on how microbes cope with the threat of cheating (Travisano & Velicer, 2004; Strassmann & Queller, 2011; Özkaya et al. 2017; Wechsler et al. 2019; Smith & Schuster, 2019). While interesting in its own right, the focus on cheating might have diverted attention from other factors that could also influence the evolution of cooperative traits (see Zhang & Rainey, 2013 for a critic).

For example, we currently know little about environments that select for increased cooperation. Additionally, it is currently unclear whether cooperative traits can be lost for reasons other than cheating, for example in environments where cooperation is less beneficial or not needed at all (Velicer et al. 1998; Zhang & Rainey, 2013).

Here, we tackle these questions by focussing on the evolution of pyoverdine production in the bacterium *Pseudomonas aeruginosa*, one of the most widely studied social traits in microbes (Griffin et al. 2004; Buckling et al. 2007; Dumas & Kümmerli, 2012; Harrison, 2013; Ross-Gillespie et al. 2015; Harrison et al. 2017; O’Brien et al. 2017). Pyoverdine is a siderophore that chelates environmental or host-bound ferric iron, and is secreted upon sensing iron scarcity. Iron-loaded pyoverdine is imported by bacteria via a specific receptor. Import is followed by iron reduction, release from the siderophore, and subsequent recycling of pyoverdine (Kramer et al. 2020; Schalk et al. 2020). Pyoverdine production is a cooperative trait, because the molecules are costly for the individual cell to produce, but once loaded with iron they become available to other cells in the local neighbourhood (Buckling et al. 2007; Harrison, 2013).

It is well established that pyoverdine, as a so-called “public good”, can select for cheating in severely iron-limited and well-mixed environments, where cheats can freely access secreted pyoverdines (Griffin et al. 2004; Kümmerli et al. 2015). It is less clear, however, how pyoverdine production would evolve in other environments where, for example, iron is less stringently limited and/or where environmental viscosity would limit cell mixing and thus cheating. Several scenarios can be envisaged. First, in iron-rich environments pyoverdine production might be selectively lost because it is not required (Zhang & Rainey 2013). Second, in viscous environments increased levels of pyoverdine production could be favoured because viscosity reduces cell dispersal and siderophore diffusion (Kümmerli et al. 2009; Julou et al. 2013), which ensures that pyoverdine-mediated social interactions occur more often between genetically related individuals. Such interactions are expected to favour cooperation (Dobay et al. 2014). Third, iron availability and environmental viscosity might interact and jointly affect the cost and benefit of pyoverdine production, and thereby select for an altered production level that matches the optimal cost-to-benefit ratio of the environment that bacteria evolved in. Finally, the social trait might diversify without the production levels being affected. Indeed, a great diversity of pyoverdine variants exists among *Pseudomonas* strains (Butaite et al. 2017), but each strain produces only one type of pyoverdine, together with its cognate receptor. Mathematical models suggest that evolutionary changes in pyoverdine and receptor structure could help individuals to escape cheating (Lee et al. 2012) or to gain an edge over competitors in the race for iron, even under iron rich conditions (Niehus et al. 2017).

To test for these alternative evolutionary outcomes, we experimentally evolved replicated populations, of the laboratory strain *P. aeruginosa* PAO1, for 200 days (approximately 1200 generations) in nine different environments. We manipulated environmental conditions along two gradients, iron availability and media viscosity, with each gradient entailing three levels in a 3×3 full-factorial design. After experimental evolution, we quantified changes in pyoverdine production and investigated the extent to which pyoverdine remained shareable among cells. We then assessed how phenotypic changes affect the fitness of evolved populations and clones. Finally, we sequenced the genomes of 119 evolved clones to map evolved phenotypes to genotypic changes.

## Material and Methods

### Strains and culturing conditions

We used the *P. aeruginosa* wildtype strain PAO1 (ATCC15692) and a fluorescent variant (PAO1-*gfp*), directly derived from the wildtype strain. PAO1-*gfp* constitutively expresses GFP (green fluorescent protein), due to a single copy insertion of *gfp* in the bacterium’s chromosome (*att*Tn7::p*tac-gfp*). We further used the pyoverdine negative mutant PAO1*ΔpvdD* (henceforth named pyoverdine-null) and the pyoverdine-pyochelin double-negative mutant PAO1*ΔpvdDΔpchEF* (henceforth named siderophore-null) as controls. The siderophore-null mutant was used because pyochelin, a secondary siderophore of *P. aeruginosa*, can partly compensate for the lack of pyoverdine (Ross-Gillespie et al. 2015).

We grew overnight pre-cultures in 8 mL Lysogeny Broth (LB - 2% m/v) and incubated at 37°C, 210 rpm, for 16-18 hours. We performed all experiments in liquid CAA medium (5 g/L casamino acids, 1.18 g/L K_2_HPO_4_.3H_2_O, 0.25 g/L MgSO_4_.7H_2_O, 25 mM HEPES buffer), to which we added 400 µM of the iron chelator 2,2′-Bipyridyl to bind the residual iron present in the medium. For the experimental evolution and follow-up experiments, we manipulated iron availability of the CAA medium by adding no FeCl_3_, 2 µM FeCl_3_ or 20 µM of FeCl_3_ to achieve conditions of low, intermediate and high iron availability, respectively. We further manipulated the viscosity of the CAA medium by adding either no, 0.1% or 0.2% (m/V) agar. Increased environmental viscosity reduces the mobility of bacteria and the diffusion of secreted compounds (Kümmerli et al. 2009). In other words, it is an experimental way to enhance spatial structure as it reduces the range across which social interactions can take place and thus increases relatedness between interacting individuals. All media components were purchased from Sigma-Aldrich (Buchs SG, Switzerland). We conducted all experiments in 96-well plates, in 200 µL of media, at 37°C and 170 rpm shaking conditions.

### Experimental evolution

We experimentally evolved strains PAO1 and PAO1-*gfp* in nine different environments, in a 3×3 full factorial design, combining the iron concentrations with the three environmental viscosities. Prior to experimental evolution, we grew overnight pre-cultures of PAO1 and PAO1-*gfp*. From these overnight cultures, we prepared glycerol stocks (750 µL culture in 750 µL glycerol (85% V/V)), stored at -80°C, to be used later as references for the ancestral strain states. We washed the remaining part of the pre-cultures in 0.85% NaCl and adjusted them to OD_600_ = 10^-2^ in NaCl (0.85% m/V), from which we inoculated 2 µL of culture into 198 µL of medium, so that experimental populations started at OD_600_ = 10^-4^ (*circa* 20000 cells/well). We evolved 24 replicates in each of our nine environments (12 populations each for PAO1 and PAO1-*gfp*, respectively), resulting in a total of 216 independently evolving populations. We inoculated the 24 replicated populations in different wells on the same 96-well plate. To reduce the risk of cross-contamination, we kept all wells adjacent to an evolving population blank. Additionally, we inoculated PAO1 and PAO1-*gfp* strains in a checkerboard pattern, so that the closest vertical and horizontal neighbors could be distinguished based on the GFP marker. We let each bacterial culture grow for 48 hours and then transferred a sample to fresh medium. We repeated this procedure for 100 consecutive growth rounds, corresponding to approximately 1200 bacterial generations. At the end of each round, we used a multimode plate reader (Infinite M-200, Tecan, Switzerland) to measure: (i) culture growth at OD_600_, (ii) pyoverdine fluorescence (excitation|emission = 400|460 nm) and, (iii) gfp fluorescence (excitation|emission = 488|520 nm). Subsequently, we diluted 2 µL of each population in 198 µL NaCl (0.85% m/V), and then transferred 2 µL of this diluted culture to 198 µL of fresh medium to begin the next round of growth, resulting in a 1:10000 dilution rate. After every other round, we mixed 50 µL of each population with 50 µL glycerol (85% V/V) for long-term storage at -80 °C.

### Cross contamination tests

We tested for potential cross-contamination events during the experiment in three ways. First, we used the constitutively expressed GFP marker to identify (i) populations that were initially untagged but started displaying GFP fluorescence beyond background-levels, and (ii) initially GFP-tagged populations that lost this trait. Second, we used a BD LSR II Fortessa flow cytometer (at the Flow Cytometry Facility, University of Zurich) to identify contaminated populations consisting of a mix of *gfp*-tagged and untagged cells. Third, we plated each population, appropriately diluted, onto LB agar plates, and screened colonies with a fluorescence imaging device (Lumenera Infinity 3 camera connected to a dark chamber) to identify *gfp*-tagged cells in untagged populations and vice-versa. We then integrated information from the three tests and excluded populations that showed signs of contamination and excluded 45 out of the 216 populations. An additional two populations were lost because they went extinct during the experiment. The maximum number of excluded populations for a given medium was eight (low iron / low viscosity). To maintain a balanced design, we randomly excluded populations from every environment with fewer than eight contaminated or extinct populations to ensure an equal number of 16 independently evolved populations per environment (i.e. a total of 144 populations) for further analysis (Table S1).

### Pyoverdine production assay

To quantify pyoverdine production changes during experimental evolution, we isolated and screened a total of 2880 evolved clones (i.e., 20 per population) for their pyoverdine production profiles. We first inoculated aliquots (from freezer stocks) of all 144 evolved populations from the last round of experimental evolution in 200 µL of LB in 96-well plates. We also inoculated the ancestral wild type strains and the pyoverdine-null mutant as controls. We grew all cultures for 16-18h, at 37°C and 170 rpm. Subsequently, we serially diluted each culture in 0.85% NaCl, to 10^-6^ and 10^-7^, and spread 50 µL of these diluted cultures onto iron-supplemented LB-Agar plates (2% m/V LB, 1.2% m/V Agar, 20 µM FeCl_3_). We incubated plates for 16-18 hours at 37°C. Subsequently, we picked 20 random colonies from each evolved population and transferred them on 96-well plates to 200 µL of the medium in which the selected clones had evolved. We further picked four colonies for each of the two ancestors (PAO1 and PAO1*-gfp*), and the pyoverdine-null mutant strains per 96-well plate. We incubated the resulting plates at 37°C and 170 rpm for 48 hours, and subsequently measured OD_600_ and pyoverdine fluorescence, using the plate reader as described above. We then quantified the evolved pyoverdine level of each clone relative to its ancestor (in % of ancestral production) and allocated evolved clones to four categories. These categories comprise: (i) non-producers, producing less than 10% of the ancestral level; (ii) reduced producers, producing between 10% and 70%; (iii) regular producers, producing between 70 % and 130 %; and (iv) hyper-producers, producing more than 130% (Dumas and Kümmerli 2012). Following our measurements, we preserved evolved clones individually by mixing 50 µL of a culture containing cells from the clone with 50 µL 85% glycerol for storage at -80°C.

### Pyoverdine growth stimulation assay

One evolutionary response to the emergence of cheaters or other competing lineages could involve changes to the peptide backbone of pyoverdines, together with mutations in receptors, such that the altered pyoverdines would become more exclusively available for members of the respective producer lineage and less exploitable by cheaters or competitors (Lee et al. 2012; Inglis et al. 2016) To explore this possibility, we used the siderophore-null mutant as a bio-sensor (Jiricny et al. 2010), and examined how well this mutant can use the pyoverdines secreted by evolved producers. Specifically, we fed the pyoverdine-containing supernatants of evolved regular producer and hyper-producer clones to the ancestral pyoverdine-null mutant and measured its growth in low iron - low viscosity condition. For this assay, we used six evolved clones from each of the 16 populations per medium (i.e. 864 clones in total). Whenever clones within a population produced similar amounts of pyoverdine, we chose six random clones. In contrast, whenever clones from the same population produced different amounts of pyoverdine (i.e. regular and hyper-producers), we chose at least one clone per pyoverdine production phenotype and drew the remaining clones at random.

To generate pyoverdine-enriched supernatants, we individually grew the selected 864 evolved clones in 200 µL of the low iron – low viscosity condition in 96-well plates, and incubated the plates for 24 hours at 37°C and 170 rpm. Following incubation, we first measured the OD_600_ and pyoverdine fluorescence of all cultures, and then separated the supernatant from the bacterial cells by centrifugation at 2250 rpm for 10 minutes. We filter-sterilized the supernatants by transferring them to 96-well 1 mL filter-plates (VWR International, Switzerland), followed by centrifugation at 1200 rpm for 15 minutes. We then performed a supernatant cross-feeding assay to quantify to extent to which each supernatant stimulates the growth of the ancestral pyoverdine-null mutant. Specifically, we added 20 µL of each supernatant, in triplicate, to individual wells of a 96-well plate containing 178 µL of the low iron – low viscosity medium. To this mixture, we added 2 µL of the siderophore-null mutant from an overnight LB culture, washed and re-suspended in 0.85% NaCl, and diluted to a starting density of OD_600_ =10^-3^. We incubated plates at 37°C and 170 rpm, and measured the growth (OD_600_) of the siderophore-null mutant after 24 hours. As positive and negative controls, we also fed the siderophore-null mutant with either supernatants of the ancestral strain or with its own supernatant. For each evolved pyoverdine producer, we calculated the cross-feeding effect as the growth (final OD_600_) of the pyoverdine-null mutant in the evolved supernatant divided by its growth in the ancestral supernatant. Values above and below one indicate increased and decreased growth stimulation, respectively.

### Sequencing of evolved clones

We sequenced the genomes of 129 evolved clones to identify the genetic changes that occurred during experimental evolution. For each of our nine growth conditions, we chose clones from 8 out of the 16 independently evolving populations. From each population, we picked 1-3 clones so that we sequenced 14 to 15 clones per condition. We chose these clones according to their evolved pyoverdine production levels, and according to how well they stimulated the ancestral pyoverdine-null mutant. Specifically, we first chose evolved pyoverdine producers with the two highest and two lowest growth stimulatory effects on the ancestral pyoverdine-null mutant (4 clones per medium = 36 clones). Second, we selected additional clones based on their evolved pyoverdine production phenotypes and growth profiles. For each population in which evolved pyoverdine phenotypes were uniform across all 20 isolated clones, we only picked a single clone for sequencing. Conversely, when pyoverdine production phenotypes varied within a population, we chose two to three clones covering the observed phenotypic range (i.e. from non- to hyper-producers). Additionally, we also sequenced one clone from each of the two ancestral strains (PAO1 and PAO1-*gfp*) as references, against which we mapped the evolved clones to identify genetic changes that occurred during experimental evolution.

Prior to DNA extraction, we grew all selected clones in 1.5 mL LB, in 24-well plates, at 37°C (170 rpm) for 16-18 hours. We extracted genomic DNA from 400 µL of these cultures with the Maxwell® RSC Cultured Cells DNA Kits using a Maxwell® RSC Instrument (Promega, Switzerland), at the Functional Genomics Centre Zurich (FGCZ). We followed the manufacturer’s instructions, including the addition of 10 µL RNase A solution to each overnight culture, to remove co-purified RNA molecules. We quantified the extracted DNA concentration with the Quantifluor dsDNA sample kit (Promega, Switzerland) and diluted to 14-32 ng/µL, when necessary. We sent 60 µL of each DNA sample to the Oxford Genomics Centre (Oxford, United Kingdom) for library preparation and whole-genome sequencing on the Illumina NovaSeq6000 platform (paired-end, 150 base-pair reads). We had to exclude 10 clones from further analysis, because we i) detected DNA cross-contaminations of samples (4 clones); ii) did not manage to extract enough genomic DNA (1 clone)); or iii) did not obtain sufficient read numbers (5 clones), which brought the final tally to 119 evolved genomes.

### Genomic analysis of evolved clones

We assessed the quality of the raw reads with “fastqc” (Andrews 2010) and “multiqc” (Ewels et al. 2016). We then aligned the reads to the *P. aeruginosa* PAO1 reference genome using the “bwa-mem” algorithm with default settings (Li 2013), followed by variant-calling with “bcftools” (Li et al. 2009): “mpileup” and “call”. We then quality-filtered the detected variants: we excluded SNPs and indels under 50 “QUAL”, as well as indels with IDV < 10. Subsequently, we excluded variants detected in the genomes of evolved clones that were also present in the ancestral clones with “bcftools isec”. We then annotated the final list of variants with “snpEff” (Cingolani et al. 2012). We detected deletions and duplications using CLC Genomic Workbench (version 11.0.1, https://digitalinsights.qiagen.com). After creating a Mapping Graph Track, we identified graph threshold areas (window size = 5) using an upper coverage threshold of 0.5 for deletions and a lower coverage threshold of 500 for duplications. We subsequently visually confirmed these results with the Integrated Genomics Viewer (Robinson et al. 2011). We identified 314 loci that independently mutated in two or more populations and classified them according to their biological function using PseudoCAP (Winsor et al. 2005) and the primary literature.

### Statistical Analysis

We performed all statistical analyses with R version 3.6.3. (R Core Team 2020). If not indicated otherwise, we used general linear models for data analysis. Prior to analysis, we used (i) the Shapiro-Wilk test and QQ-plots to check whether the model residuals were normally distributed; and (ii) the Levene test to test for homogeneity of variances. If these assumptions were not met, we log-transformed the affected response variables.

We first tested whether media conditions affected end-point growth (OD_600_ after 48 hours) and pyoverdine production of the ancestral strain. In our linear models, we fitted either of these two population-level metrics as dependent variables, and iron availability and media viscosity as response variables. We quantified pyoverdine investment as “PVD per capita”, i.e. the pyoverdine fluorescence normalized by OD_600_, and logarithmically transformed these values.

To determine whether pyoverdine investment changed between the first and the last round of experimental evolution, we calculated the fold change in pyoverdine investment for each population at round 100 relative to round 1, and performed one-tailed t-tests to evaluate whether the observed values differ from the null expectation (i.e. fold-change = 1). We used the same procedure to assess how the supernatant of evolved clones changed the growth of the ancestral non-producer relative to the supernatant of the ancestral producer. We used the FDR p-value adjustment method to account for multiple testing (Benjamini & Hochberg, 1995).

To test whether evolved populations harbour different coexisting pyoverdine phenotypes, and whether the heterogeneity in phenotypes differs across environments, we: (i) quantified the coefficient of variation for each population (standard deviation/mean across 20 clones) and (ii) then fitted this metric as a dependent variable, together with media viscosity and iron availability as response variables to a linear model. We logarithmically transformed coefficients of variation for statistical analysis and used Tukey’s *post hoc* tests to examine pair-wise differences between factor levels.

To test whether the frequency of evolved pyoverdine non-producers and over-producers differ across the nine experimental media, we used Fisher’s exact test, with FDR p-value corrections for multiple testing. We used the same statistical test to evaluate whether hypermutator and non-hypermutators clones differ in the number of mutations in pyoverdine biosynthesis versus regulatory genes.

To test whether clones from the four pyoverdine production categories differ in their growth, we quantified the OD_600_ of the evolved clones after 48 hours and scaled all values relative to the ancestral producer. We then fitted this metric as a response variable in linear mixed models (lme4 package: Bates et al. 2015), and used category as a response variable, and population of origin as a random factor. Finally, to assess whether mutations in functional categories are associated with changes in pyoverdine production and growth, we fitted linear models where we used pyoverdine production relative to the ancestor as the response variable, and absence/presence of mutations in the respective category as the explanatory variable. Since mutations in the pyoverdine locus had a significant effect on pyoverdine production, we always implemented the absence/presence of mutations in this locus as an additional explanatory variable in our models.

## Results

### Interaction between iron availability and environmental viscosity determine pyoverdine production and growth in ancestral P. aeruginosa

We first evaluated how iron availability and media viscosity affect the growth and pyoverdine production of the ancestral *P. aeruginosa* prior to evolution (Figure 1). This analysis provides important information on the extent to which phenotypes differ across environments and thus the potential for selection to act differentially upon them. We found that ancestral *P. aeruginosa* growth was influenced by a significant interaction between iron availability and media viscosity (ANOVA: F_4,135_ = 55.43; p < 0.0001, Fig. 1A, Table S2). While growth decreased with more stringent levels of iron limitation, increased media viscosity dampened this effect. Pyoverdine production was also influenced by a significant interaction between iron availability and media viscosity (ANOVA: F_4,135_ = 237.67; p < 0.0001, Fig. 1B, Table S2). While pyoverdine production gradually decreased from low to high iron availability, the decrease was again less pronounced in more viscous environments. These findings show that *P. aeruginosa* adjusts its pyoverdine production profile both in response to iron availability and to environmental viscosity, and we might thus expect selection pressures on this trait to vary across environments.

**Figure 1.**
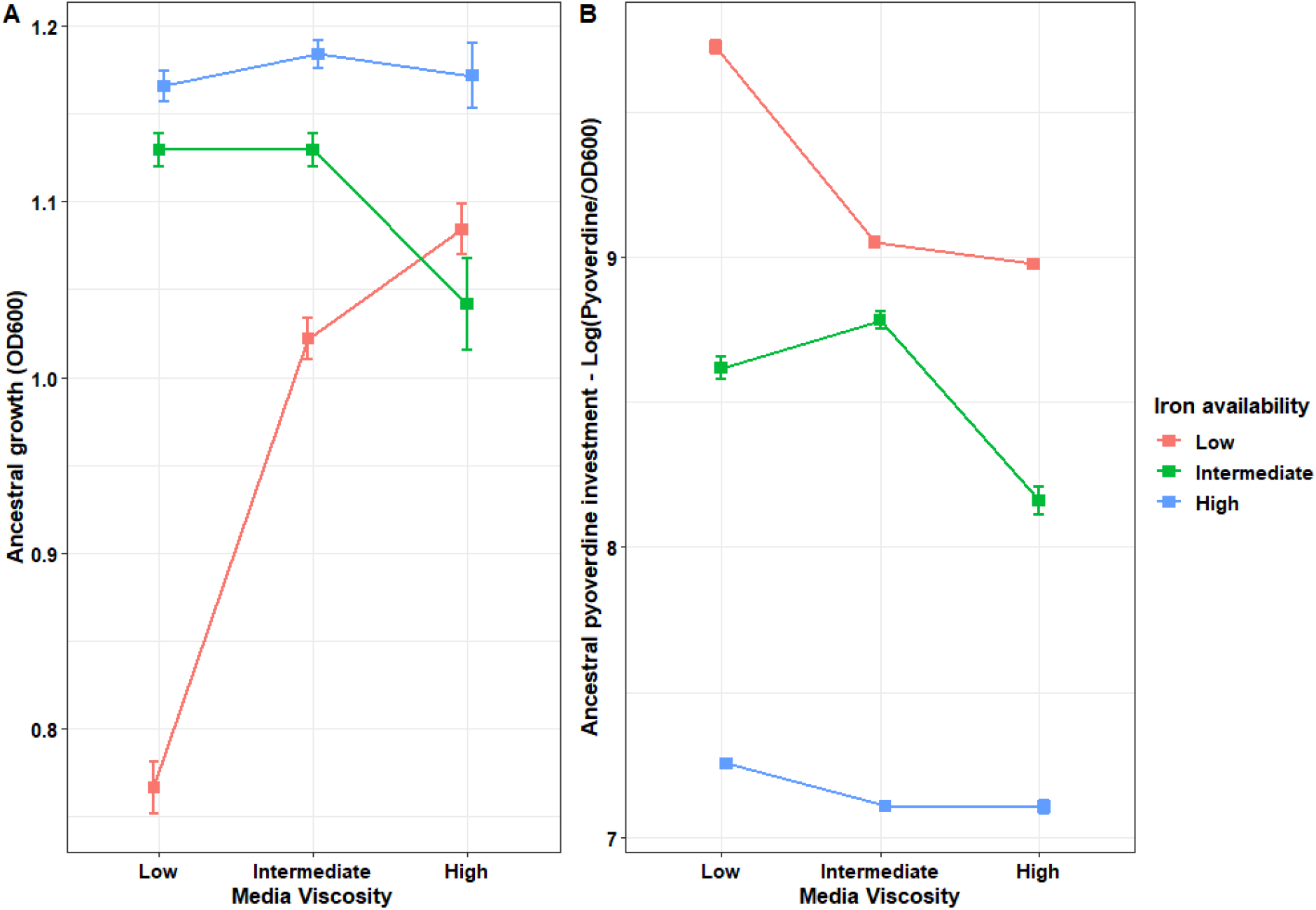
Iron availability and media viscosity interact and jointly influence growth and pyoverdine production of the *P. aeruginosa* PAO1 ancestral strain. (A) Strain growth (OD at 600 nm) in the nine different environments, which vary in medium viscosity (amount of agar added: low = 0.0%; intermediate = 0.1%, high = 0.2%) and iron availability (low = 0 µM, red line; intermediate = 2 µM, green line; high = 20 µM, blue line). (B) Pyoverdine investment, shown as log-transformed pyoverdine fluorescence values normalized by OD, in all nine experimental conditions. Each square represents the mean of 16 independent populations, whereas the bars represent the standard error of the mean.

### Environment-dependent selection for decreased and increased pyoverdine production

To test whether different environments select for different pyoverdine production levels, we quantified the pyoverdine investment of evolved populations relative to the ancestor after approximately 1200 generations of experimental evolution. We observed that pyoverdine production levels significantly changed in five out of the nine environments (Figure 2A). Consistent with previous studies, we found that populations evolved reduced pyoverdine production levels under the low iron – low viscosity condition (t_15_ = -2.617, p_adj_ = 0.035). Because iron limitation is most stringent and pyoverdine most needed in this environment (Fig. 1), the evolution towards lower pyoverdine production levels is compatible with the arise and spread of cheating mutants. In stark contrast, populations that evolved in low viscosity environments but with intermediate and high iron availabilities significantly increased pyoverdine production levels compared to the ancestral wildtype (intermediate iron – low viscosity: t_15_ =7.134, p_adj_ < 0.0001; high iron – low viscosity: t_15_ = 5.830, p_adj_ < 0.0001).

**Figure 2.**
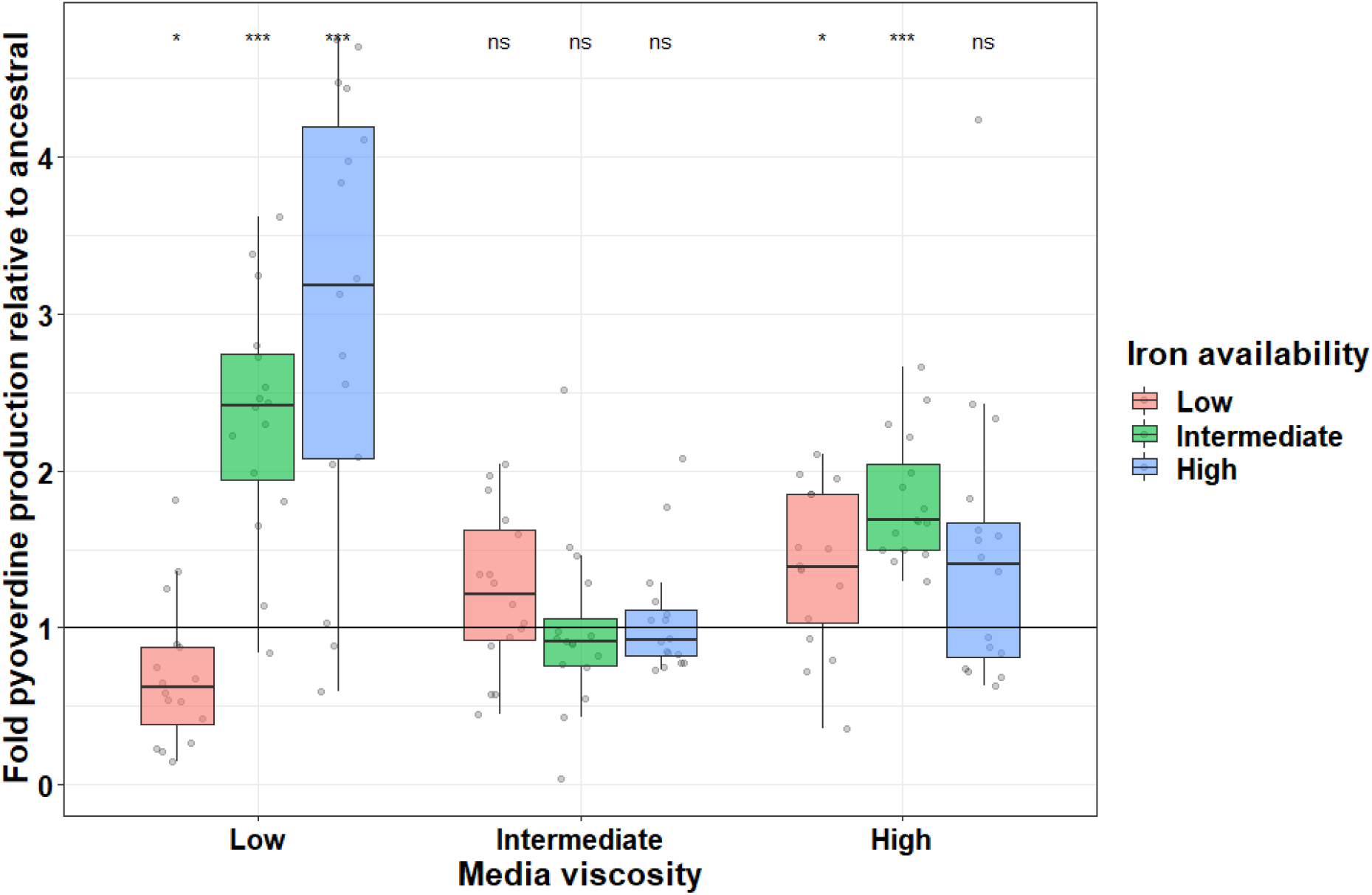
Evolution causes a media-dependent divergence in pyoverdine production. Population-level pyoverdine production after 100 rounds of experimental evolution, relative to the ancestral phenotype (black horizontal line) in nine different environments. The environments varied in their viscosity (amount of agar added: low = 0.0%; intermediate = 0.1%, high = 0.2%) and iron availability (low = 0µM, red; intermediate = 2 µM, green; high = 20 µM, blue). We measured pyoverdine production via its natural fluorescence, scaled to the optical density of the population (OD at 600 nm after 48 hours). Box plots show the median and the 1^st^ and 3^rd^ quartile across the 16 independently evolved populations (depicted by the individual dots). Whiskers represent the 1.5 interquartile range. Asterisks signify FDR-corrected p-values for one-sample t-tests: * < 0.05, *** p < 0.0001, ns = not significant.

Selection for altered pyoverdine production seemed to be lowest in media with intermediate viscosity, where the population-level pyoverdine production levels did not change during experimental evolution regardless of iron concentration (intermediate viscosity with low iron: t_15_ = 1.865, p_adj_ = 0.105; intermediate iron: t_15_ = −0.134, p_adj_ = 0.895; high iron: t_15_ = 0.584, p_adj_ = 0.639). Conversely, in populations evolved in high viscosity media, we observed an increase in pyoverdine production levels in most populations. This increase was significant for two out of the three iron conditions (high viscosity - with low iron: t_15_ = 2.985, p_adj_ = 0.021; with intermediate iron: t_15_ = 8.186, p_adj_ < 0.0001; with high iron: t_15_ = 2.112, p_adj_ = 0.078). Taken together, our results do not support the hypothesis that pyoverdine production is selected against in iron-rich environments because of potential disuse. Instead, we found support for the hypothesis that high environmental viscosity can favour increased levels of cooperation and that pyoverdine production levels diverge across environments, possibly to match environment-specific requirements for this siderophore.

### Iron availability and viscosity both influence heterogeneity in pyoverdine production among clones from the same population

In a next step, we studied evolved populations individually and asked whether clones with distinct pyoverdine phenotypes co-exist within populations. We predict highest population diversity in environments where different social strategies are likely to compete, e.g. under low iron – low viscosity, where cheats and pyoverdine producers could closely interact. Conversely, we expect lower levels of diversity among clones in environments that are generally favourable for cooperation (e.g. under high viscosity). Indeed, we observed that the level of within-population heterogeneity in pyoverdine production varied both between environments and between replicates of the same condition (Fig. S1). In line with our hypothesis, we found that the coefficient of variation, which quantifies phenotypic heterogeneity among clones within a population, significantly increased with more stringent iron limitation (ANOVA: F_2,139_ = 4.85; p = 0.009; Tukey post-hoc tests: low vs intermediate t = - 2.993, p = 0.009; low vs high (marginally significant) t =-2.243, p = 0.067) and lower environmental viscosity (ANOVA: F_2,139_ = 11.34; p < 0.0001; Tukey post-hoc tests: low vs intermediate t = -3.823, p < 0.001; low vs high t = -4.37, p < 0.001) (Fig. 3A, Table S3).

**Figure 3:**
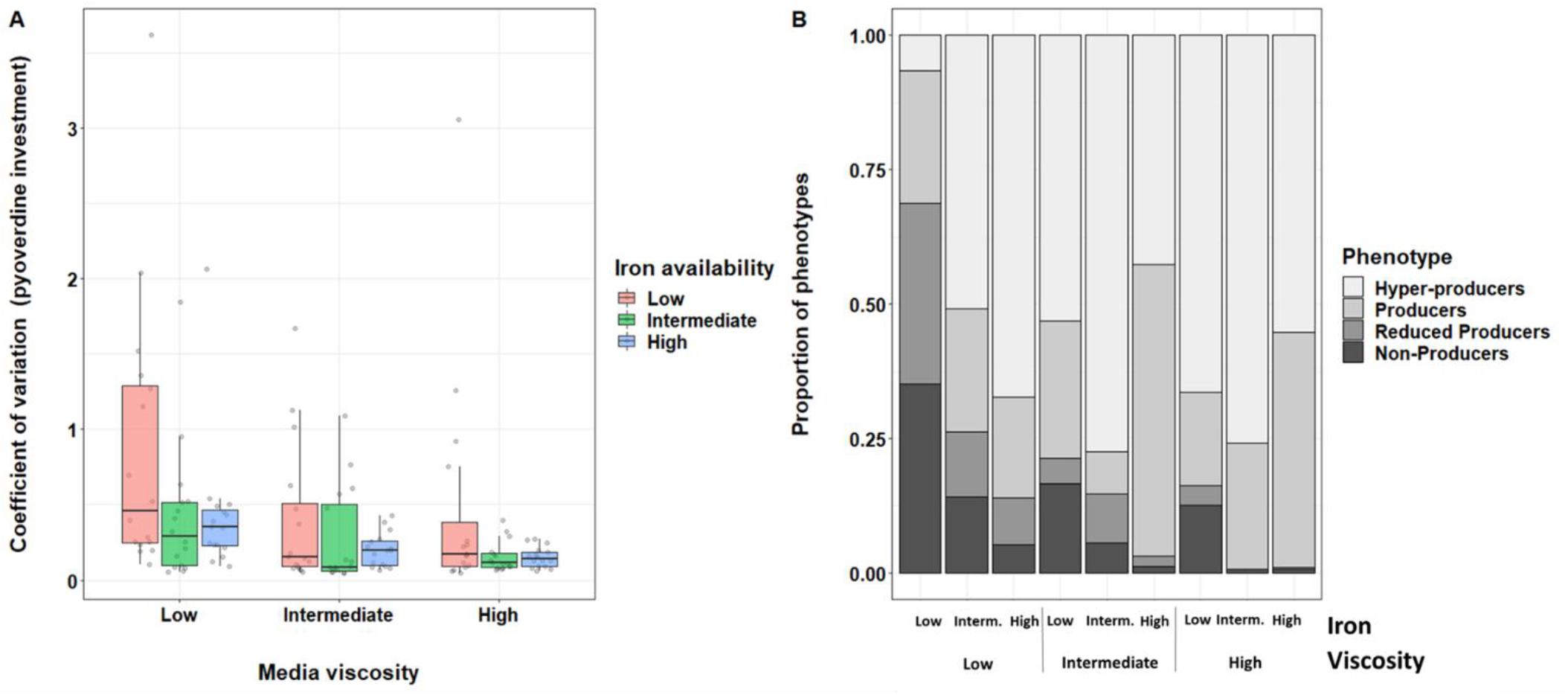
Clones with divergent pyoverdine production profiles co-exist in populations evolved under low iron or viscosity conditions, whereas populations evolved under higher iron and high viscosity consist almost exclusively of producers and hyper-producers. (A) Coefficient of variation (standard deviation/mean) for pyoverdine production phenotypes across clones from the same population. We measured pyoverdine production via its natural fluorescence and scaled to the optical density (OD at 600 nm) of the population. Box plots show the median and the 1^st^ and 3^rd^ quartile across the 16 independently evolved populations (depicted by the individual dots). (B) Distribution of discrete pyoverdine production phenotypes for all nine environments after 100 rounds of experimental evolution. We allocated clones to phenotype categories according to their pyoverdine production levels scaled relative to the ancestor (100%). [1] Non-producers: less than 10%; [2] reduced-producers: 10%-70%; [3] producers: 70 %-130%; [4] hyper-producers: more than 130% of ancestral production. Fisher’s exact test reveals significant differences in category frequencies between any two environments.

We then examined whether the different environments affect the prevalence of clones with distinct pyoverdine production levels. Consequently, we allocated each isolated clone into one of four discrete pyoverdine production categories, covering the range from non-producers to hyper-producers (Fig 3B). We found that the frequency of clones in the four categories varied significantly between any two environments (global Fisher’s exact test: p < 0.0005; FDR-corrected pairwise Fisher’s exact tests maximum: p < 0.0079). Most importantly, the frequency of non-producers decreased in environments with increased iron availability and media viscosity, whereas the prevalence of regular and hyper-producers increased in more viscous environments under all iron availabilities (Fig. 3B). This analysis thus reinforces the view that there is a shift from selection for cheating to selection for increased cooperation when moving from low iron – low viscosity to higher iron – higher viscosity conditions.

### Pyoverdine under- and hyper-production comes at a fitness cost

The high frequency of pyoverdine none- and hyper-producers suggests that this trait was under selection during experimental evolution. Here, we explore whether changes in pyoverdine phenotypes have fitness consequences for evolved clones. If non-producers evolved because of cheating, then their fitness should decrease when growing in monoculture. Conversely, if over-producers evolved because higher levels of cooperation are beneficial then their fitness should exceed the fitness of regular producers. Indeed, we observed significant growth differences between evolved clones from the four pyoverdine production categories (linear mixed model: t_373_ = 51.91, p < 0.0001, Fig. 4). As predicted by the cheating hypothesis, non-producers grew significantly more poorly than regular pyoverdine producers (Tukey post-hoc test: z = 4.024; p < 0.001). However, regular pyoverdine producers also grew significantly better than hyper-producers (z = -9.45, p < 0.001), thus refuting our hypothesis that over-production is associated with an overall growth advantage.

**Figure 4:**
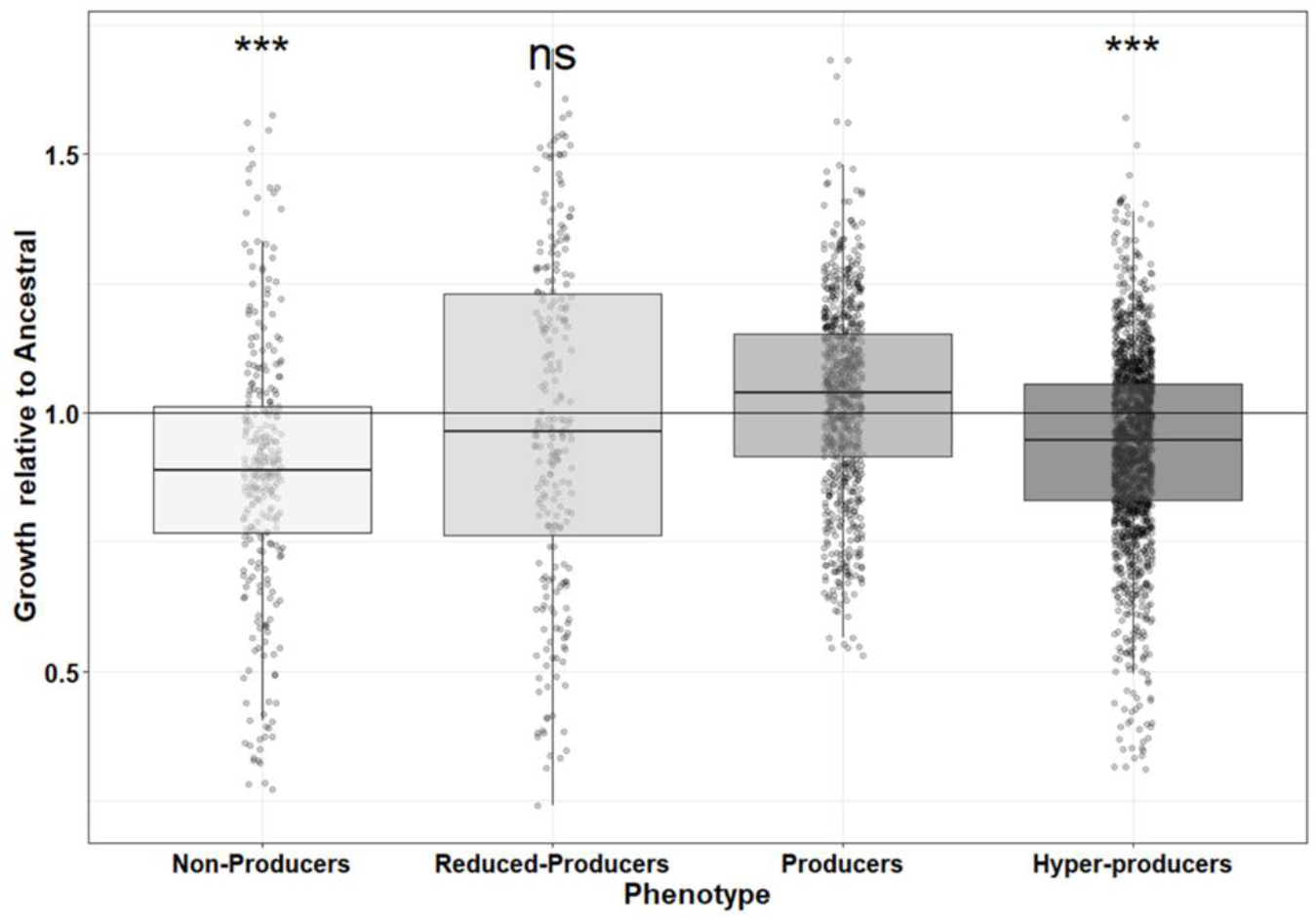
Evolution of non- and hyper-pyoverdine production is associated with growth declines. (A) Growth of evolved clones allocated to four different pyoverdine production categories, scaled relative to the ancestor (100%): None-producers: less than 10%; [2] reduced-producers: 10%-70%; [3] regular producers: 70 %-130%; [4] hyper-producers: more than 130% of ancestral production. Significance levels (*** p < 0.0001, ns = non-significant) refer to pair-wise comparisons between producers and the other three phenotype categories. Horizontal black line at y = 1 indicates the ancestral phenotype.

One shortcoming of the above analysis is that it combines clones across all environments, which does not allow us to test whether pyoverdine hyper-production is associated with fitness benefits only in some environments. To address this issue, we repeated our analysis for the nine environments separately. While we still observed that hyper-producers generally grew less well than regular producers (Fig. S2; Table S4), we also found that their growth decline was most pronounced in the low iron – low viscosity condition. In contrast, growth differences between producer and hyper-producer clones became negligible in environments with increased iron or increased viscosity, suggesting that pyoverdine hyper-production might be equally successful as regular production in these environments. Moreover, it is possible that hyper-producers show increased growth rates or shorter lag-phases, fitness components we could not measure with our end-point assay.

### Environment-dependent change in the extent to which non-producers are stimulated in their growth by pyoverdines from evolved producers

Since the above analyses showed that most pyoverdine non-producers may have evolved as cheaters, we here explore whether pyoverdine producers and hyper-producers responded to the presence of non-producers by reducing the exploitability of their pyoverdine. To test this hypothesis, we harvested pyoverdine-containing supernatants from ancestral and evolved (hyper-)producers, and fed these supernatants to an engineered siderophore-null mutant. If (hyper-)producers evolved mechanisms to reduce the exploitability of their pyoverdine, we expect their supernatants (containing pyoverdine) to have a reduced stimulatory effect on the growth of the siderophore-null mutant.

We found that the supernatants of all evolved (hyper-)producing clones still stimulated the growth of the ancestral non-producer (Fig. 5). However, the growth stimulation experienced by the siderophore-null mutant depended on a significant interaction between the viscosity and iron availability of the environments in which the (hyper-)producers evolved in (LMER: 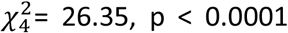, Fig. 5, Table S5). Compatible with our hypothesis, we observed that the relative growth stimulation of the siderophore-null mutant was significantly reduced when subjected to supernatants from producers evolved in the low iron – low viscosity environment (t_35_ = -6.40, p < 0.0001), exactly the environment in which non-producers were most prevalent (Fig. 3C). In all other environments, the supernatant-induced growth stimulation of the siderophore-null mutant was either unchanged or even higher compared to the supernatant of the ancestral pyoverdine producer (Fig. 5). Increased growth stimulation can be explained by the fact that hyper-producers evolved in many treatments, such that the pyoverdine content in the supernatant was an important additional predictor of the observed growth stimulation patterns (LMER: 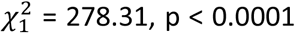, Table S5). Taken together, these results suggest that only pyoverdine (hyper-)producing clones evolved under low iron – low viscosity might have adapted to non-producers and reduced the exploitability of their pyoverdine.

**Figure 5.**
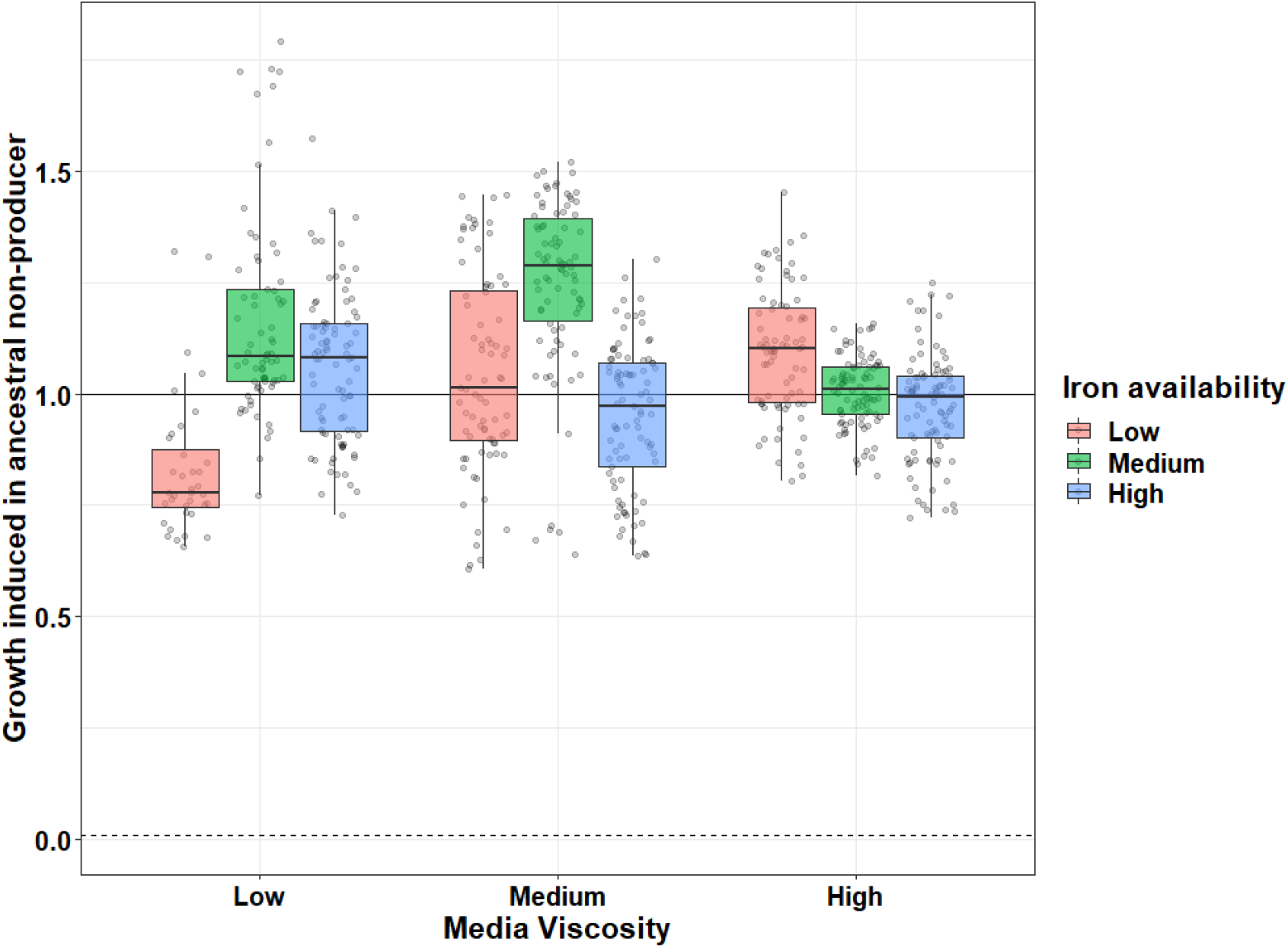
Interaction between iron availability and environmental viscosity drives the extent to which pyoverdines from evolved producers stimulate the growth of the ancestral non-producer. Each individual circle shows the average effect (three replicates) of supernatant collected from an evolved pyoverdine (hyper-)producers on the growth of an engineered siderophore-null mutant. All growth values are scaled relative to the growth stimulation this mutant experienced in the supernatant of the ancestral wildtype (solid horizontal line). The dashed line represents the growth of the siderophore-null mutant when fed with its own supernatant. Box plots show the median and the 1^st^ and 3^rd^ quartile, whereas whiskers represent the 1.5 interquartile range.

### Genomic features of evolved clones

To map the evolved phenotypes to genetic changes, we sequenced the genomes of 119 evolved clones from 68 independent populations to an average sequencing depth of 230x (min = 177x, max = 334x), and compared them to their respective ancestors (PAO1 and PAO1-*gfp*). We found a total of 3472 mutations (SNPs, duplications and deletions), distributed over 1897 loci (coding and intergenic regions). Individual clones had between 2 and 191 mutations (median = 7, Fig. 6A). The distribution of mutations per clone followed a bimodal pattern: most clones (78.99%) had few mutations (< 20), while a minority of clones (21.01%) had many mutations. This discrepancy in mutations per clone can be explained by the occurrence of non-synonymous mutations in the coding regions of the *mutS* and *mutL* genes in almost all clones (92%, 23 out of 25) possessing more than 20 mutations (Fig. 6A). These genes code for parts of the DNA mismatch repair system of *P. aeruginosa*, and mutations in their coding regions result in substantially elevated mutation rates (Oliver et al. 2002; Wiegand et al. 2008). We henceforth call clones harboring such mutations hyper-mutators.

**Figure 6:**
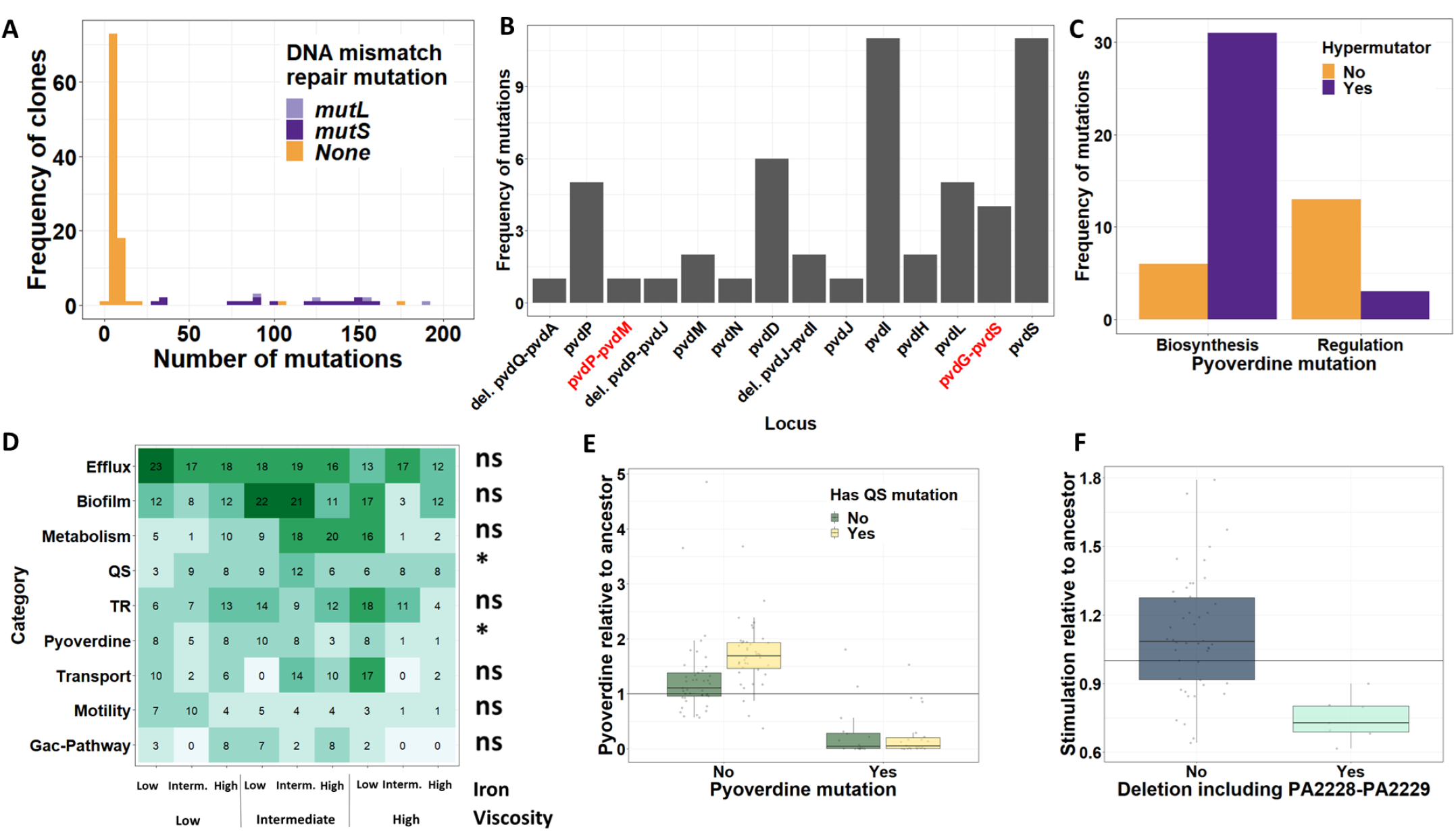
Whole-genome sequencing reveals that mutations in the pyoverdine locus and quorum sensing genes are respectively responsible for pyoverdine under- and hyper-production. (A) Distribution of the number of mutations per clone. All but two clones with more than 20 mutations have mutations in the DNA mismatch repair machinery and are therefore considered hypermutators. (B) Distribution of mutations in genes (black labels) and intergenic regions (red labels) of the pyoverdine locus across clones. Genes are ordered according to the physical map of the locus. (C) Non-hypermutator clones have more mutations in pyoverdine regulatory genes, whereas mutations in pyoverdine biosynthesis genes are over-represented among hypermutator clones. (D) Heatmap of the most common mutational targets sorted by functional categories, across our nine different environments. TR – Transcriptional regulation, QS – Quorum sensing. Labels to the right of the heatmap indicate whether mutations in the respective category had a significant (*, p < 0.05) or non-significant (ns, p > 0.05) effect on evolved pyoverdine production level. (E) Box plots showing the evolved level of pyoverdine production for clones with mutations in the pyoverdine locus, in QS genes, in both types of genes, or in none of these genes. (F) Box plots showing the evolved level of growth stimulation of the ancestral non-producer, for clones that have or do not have a deletion that includes the upstream region and/or part of the qsrO-vqsM-PA2228 operon. Box plots in (E) and (F) depict the median, the 1^st^ and 3^rd^ quartile and the 1.5 interquartile range (whiskers). The horizontal black lines at y = 1 indicate the ancestral phenotypes.

### Mutations in the pyoverdine locus are associated with reduced pyoverdine production

We then focused on non-synonymous mutations in the pyoverdine locus, i.e., in the regulatory and biosynthetic genes for pyoverdine production, together with their intergenic regions. We asked whether mutations in these genes affect the pyoverdine production of evolved clones. A total of 38 clones had mutations in the pyoverdine locus, and these mutants produced significantly less pyoverdine (mean: 0.27 ± SE: 0.07) than clones without such mutations (mean: 1.50 ± SE: 0.08) (ANOVA: F_1,117_ = 88.86; p < 0.0001, Fig. S3). Only one clone produced no pyoverdine without having a mutation in the pyoverdine locus. We could attribute its phenotype to a non-synonymous mutation in *tatC*. Mutations in this gene are known to impair pyoverdine maturation (Lee et al. 2016; Voulhoux et al. 2006), and we thus included this mutant in the class of pyoverdine mutants.

Almost all genes of the pyoverdine locus harbored mutations (Fig. 6B). The most frequent mutations occurred in or upstream of *pvdS*, the sigma factor that positively regulates the expression of pyoverdine, and in *pvdI*, the largest gene in the cluster, encoding a non-ribosomal peptide synthetase (Fig. 6B). While *pvdS* mutations have frequently surfaced in previous more short-term evolution experiments (Granato et al. 2018; Kümmerli et al. 2015), mutations in pyoverdine biosynthesis genes are much less common (O’Brien et al. 2019). We suspected that this difference is linked to the evolution of hypermutators in our experiment. Indeed, hypermutators incurred significantly fewer mutations in regulatory genes, but more mutations in pyoverdine biosynthesis genes compared to non-hypermutators (Fisher’s Exact Test: p < 0.0001, Fig. 6C). For all those populations where we sequenced more than one clone, we examined the order in which mutations appeared (Fig. S4). We found that mutations in the pyoverdine locus arose after *mutS*/*mutL* mutations in eight populations, but never the other way round. In six populations, we were not able to resolve the order in which mutations occurred. Taken together, we observed a high prevalence of mutations in the pyoverdine locus, a shift in the mutational landscape between non- and hypermutators, and most mutations to be associated with decreased pyoverdine production.

### Mutations in quorum-sensing systems are associated with pyoverdine hyper-production

Next, we investigated possible links between mutational patterns and pyoverdine hyper-production. For this purpose, we classified mutations into functional categories. Fig. 6D shows the nine most frequent categories (in which we detected at least 25 independent mutations per category), and the prevalence of mutations in them across our nine environments. One of these categories (“pyoverdine”) includes mutations in the pyoverdine locus itself, which we have shown to be associated with under-production. We thus asked whether mutations in any of the other eight categories are associated with pyoverdine hyper-production, while statistically controlling for the presence of mutations in the pyoverdine locus. We found that mutations in quorum-sensing (QS) genes were significantly associated with increased pyoverdine production in clones with no mutations in the pyoverdine locus (ANOVA: F_1:115_ = 8.211, p_adj_ = 0.039) (Fig. 6E; Table S6). Mutations occurred in all three known *P. aeruginosa* QS systems (Las, Rhl, and PQS) and included non-synonymous SNPs, small indels, and large duplications and deletions (Table S7). In addition, the mutational patterns of Fig. 6D also suggest that some mutations are associated with adaptation to the growth medium. The most obvious candidates are mutations in efflux pump related genes, particularly those responsible for the regulation of the mexAB-oprM pump (Table S8), which occurred in 108 out of the 119 clones. Their high prevalence could result from the use of bipyridyl, a compound that not only chelates iron outside the cell, but has also toxic effects when entering cells. Efflux pump up-regulation has been reported to improve growth in the presence of bipyridyl (Liu et al. 2010).

### Deletions in the qsrO-vqsM-PA2228 operon and its upstream region are associated with a reduction in pyoverdine-mediated growth stimulation

Finally, we investigated whether the reduction in pyoverdine-mediated growth stimulation observed for some evolved producer clones (Fig. 5) is associated with specific genetic changes. We followed a similar strategy as above (Fig. 6D), testing whether mutations in genes from specific functional categories are associated with a reduction in pyoverdine-mediated growth stimulation. This approach yielded no significant association for any category (Table S9). We therefore explored whether mutations in specific loci were associated with changes in this phenotype. We found a single hit: supernatants from clones with a large deletion (130 bp to 21 kb) comprising the *qsrO-vqsM-*PA2228 operon and its upstream region (7 cases) stimulated the growth of the ancestral non-producer significantly less than clones without such deletions (Fig 6F, ANOVA: F_1:51_ = 13.979, p_adj_ = 0.007, Table S10).

## Discussion

Because bacterial social traits, such as biofilm formation, quorum-sensing (QS) communication, and the secretion of beneficial compounds affect virulence, eco-system functioning and microbiome assembly, enormous interest has focused on them (Magnúsdóttir et al. 2015; Davis & Isberg, 2019; Ebrahimi et al. 2019). Particular attention has been paid to cheating (Strassmann & Queller, 2011; Harrison et al. 2017; Tarnita, 2017) – the loss of cooperation through mutants that exploit the cooperative efforts of others – whereas the evolution of cooperative traits in environments that do not necessarily favour cheating remain poorly examined (but see Velicer et al. 1998). Here we explored how different environments shape the evolutionary trajectory of a model bacterial cooperative trait, the production of the iron-scavenging siderophore pyoverdine by the opportunistic pathogen *P. aeruginosa*. We experimentally evolved this species for approximately 1200 generations in nine different environments, varying in their degree of iron availability and environmental viscosity. We found that selection for reduced pyoverdine production due to cheating occurred only in one of the nine environments (low iron – low viscosity), where relatedness among individuals is low and the benefit of exploitation is high. In stark contrast, pyoverdine production either did not change or even increased over time in four environments each (Fig. 2). Selection for increased pyoverdine production occurred predominantly in environments with either high iron availability, where the social trait is expressed at a relatively low level and thus not so costly, or with high viscosity, where interacting individuals are more closely related. Whole-genome sequencing revealed that point mutations in genes of the pyoverdine locus, particularly the iron-starvation sigma factor *pvdS* are significantly associated with decreased pyoverdine production. Conversely, SNPs and large scale deletions in QS genes are associated with pyoverdine hyper-production. Together, our results suggest that bacterial social traits can undergo rapid evolutionary change and allow bacteria to adapt to the prevailing environmental and social conditions.

The evolution of pyoverdine non-producers or reduced-producers only consistently occurred in one specific environment (low iron, low viscosity). Because pyoverdine is most needed under low iron availability, our findings indicate that non-and low-producers have a selective advantage due to cheating (Fig. 2, Fig. S1). This pattern is consistent with previous studies, where non-producers spread because they used pyoverdine produced by others. In doing so, they did not contribute to the production of this public good themselves, which gave them a metabolic advantage (Ghoul et al. 2014; Kümmerli et al. 2015; Harrison et al. 2017). The mutational patterns discovered are also consistent with previous work showing that pyoverdine non- or reduced-production predominantly arose by mutations in *pvdS* and its promotor region in non-hypermutator clones (Kümmerli et al. 2015; Granato & Kümmerli, 2017; O’Brien et al. 2019; Tostado-Islas et al. 2020). The sigma factor *pvdS* regulates the expression of the entire pyoverdine biosynthesis machinery (Cunliffe et al. 1995; Ringel & Brüser, 2018). Mutations in it are thus often associated with (i) reduced production of this public good, and (ii) major cost saving when the entire pyoverdine locus is shut down. In contrast to non-hypermutators, we found that mutations in pyoverdine biosynthesis genes were overrepresented in hypermutators. We hypothesize that mutations in these genes that reduce pyoverdine production costs are rare, and thus only surface in clones with elevated mutation rates.

In our long term evolution experiment, the occurrence and spread of non-producers never lead to a “tragedy of the commons”, i.e. the fixation of cheaters and the concomitant collapse of populations. Such a scenario has been proposed for several microbial social traits (Rankin et al. 2007; Kümmerli et al. 2015; Schuster et al. 2017). One possible explanation for its absence is that most cheater clones still produced some pyoverdine, allowing them to grow even in the absence of regular pyoverdine producers (Fig. 4) (Jiricny et al. 2010). Moreover, pyoverdine secretion is not the only strategy to scavenge iron. Previous studies showed that pyoverdine non-producers can switch to alternatives, such as the production of the secondary siderophore pyochelin (Ross-Gillespie et al. 2015), and the ferric-iron reducing agent pyocyanin (O’Brien et al. 2017). Pyoverdine production can further be subject to negative-frequency dependent selection, thus permitting cheaters and cooperators to co-exist (Ross-Gillespie et al. 2007).

Besides these established mechanisms, our data also suggest that pyoverdine producers can directly adapt to the presence of cheats by reducing the benefits of (or access to) their pyoverdines (Fig. 5). We observed this phenomenon predominantly in evolved producers from the low iron / low viscosity environment, where non-producer prevalence was highest and thus producer counter-adaptation most expected. While we are unable to identify the exact molecular mechanism behind the reduced pyoverdine-mediated growth benefits for non-producers, we found one mutational target, the qsrO-vqsM-PA2228 operon and its upstream region, whose deletion is associated with this effect (Fig. 6F). The exact function of this operon is unknown, but deletions in it can lead to a complete shutdown of all three QS systems of *P. aeruginosa* (Köhler et al ., 2014; Liang et al. 2014). Given that *vqsM* is a putative transcriptional regulator (Huang et al. 2019), we believe that the change in pyoverdine benefits is not a direct consequence of QS silencing, but rather a pleiotropic effect of a yet to be described regulatory link of this operon to siderophores. In any case, our findings show that adaptations to cheating are possible but did not involve the proposed modification of pyoverdine structure and its specific receptor (Smith et al. 2005; Lee et al. 2012; Inglis et al. 2016; Niehus et al. 2017; Stilwell et al. 2018). This hypothesis has been put forth multiple times, based on the observation that many pyoverdine variants exist in nature. We argue that there were ample opportunities for pyoverdine structural changes to occur in our long-term experiment, but we observed none. This suggests that pyoverdine diversification might not be driven by cheating, but other selection pressures, such as escaping phages or toxins that use the pyoverdine receptor as an entry point (Smith et al. 2005).

Another main finding of our experiment is the evolution of pyoverdine hyper-production in multiple environments, highlighting that not only cheating but also increased cooperation can be favoured. Specifically, pyoverdine production increased in high viscosity environments. High viscosity hampers individual dispersal, which increases the probability that social interactions occur between related individuals, which is favourable for cooperation (Hamilton 1964; Queller 1994; El Mouden and Gardner 2008; Dobay et al. 2014). While previous work showed that high relatedness can maintain cooperation in short-term experiments (Griffin et al. 2004; Gilbert et al. 2007; Kümmerli et al. 2009; Bastiaans et al. 2016), we here show that it can favour increased levels of cooperation during a long-term experiment. We further observed that hyper-producers emerged and spread in low viscosity environments with intermediate and high levels of iron. This was rather unexpected, because relatedness is low in these environments, and cooperation should thus be more vulnerable to cheating. However, although non-producers indeed surfaced in these environments, they never reached high frequencies (Fig. 3B). We speculate that increased pyoverdine production could be sustainable in these environments, because the cost-to-benefit ratio of this trait is altered. Cost might be relatively low because the pyoverdine production level is still considerably lower compared to what strains produce in the low iron / low viscosity environment. Benefits might be relatively high because pyoverdine is still required to scavenge iron bound to the bipyridyl chelator.

We found that mutations in QS regulatory genes (*lasR, rhlR* and *pqsR*) and full deletions of the Las-regulon were associated with an increase in pyoverdine production, suggesting a (direct or indirect) link between pyoverdine and QS regulons. We propose two non-mutually exclusive reasons to explain why these mutations were favoured in our experiment. First, mutations in QS loci may be advantageous *per se*, and pyoverdine hyper-production may simply be a by-product of these mutations. In our growth medium, QS-controlled traits such as proteases, biosurfactants, and phenazine toxins are not needed, such that abolishing quorum sensing could save substantial metabolic costs (Özkaya et al. 2018). In support of this by-product hypothesis, we found that QS-mutants occurred in all environments (Fig. 6D), suggesting that it is generally favourable to lose QS. Second, mutations in QS genes may be advantageous because they cause pyoverdine hyper-production. Our results also support this hypothesis, because pyoverdine hyper-producers did not spread in the low iron / low viscosity environment, where cheaters dominated. Instead, they reached high frequencies in high viscosity environments, where cooperation is predicted to be favourable. Thus, mutations in the QS-regulon may accomplish two goals at one blow: the silencing of a superfluous regulon, and the possibility to increase pyoverdine cooperation.

The evolutionary trajectories and associated genetic changes we describe are remarkably similar to patterns of *P. aeruginosa* evolution during chronic infections in human patients (Marvig et al. 2015; Winstanley et al. 2016). For example, longitudinal studies on cystic fibrosis patients suffering from *P. aeruginosa* lung infections revealed that social traits, particularly quorum sensing and pyoverdine production, are under selection (Jiricny et al. 2014; Andersen et al. 2015). Especially the widespread accumulation of quorum-sensing mutants that we observed in the laboratory has parallels in human lungs (Wilder et al. 2009; Bjarnsholt et al. 2010) and in animal infections models (Granato et al. 2018). Moreover, hypermutators and isolates that over-express efflux pumps are also commonly observed in clinical isolates (Sobel et al. 2005; Mena et al. 2008; Rees et al. 2019), where they seem to contribute to antibiotic resistance. Consistently, we found hypermutators in almost all our experimental treatments, and the widespread arising and spreading of efflux pump mutants, probably in response to bipyridyl toxicity. Taken together, these parallels suggest that *in vitro* studies, despite their obvious limitations, may help understand evolutionary trajectories taken by pathogens.

In conclusion, we found that *P. aeruginosa* can quickly evolve alternative social phenotypes to match prevailing abiotic (iron availability) and biotic (relatedness) conditions. Cheating, which was the focus of many previous studies, seems to be only favored under very specific conditions, and thus plays a relatively minor role in the environments studied here and perhaps also in many other environments. Instead, our data suggest that *P. aeruginosa* adapts its pyoverdine production profile to match environmental requirements, often by up-regulating pyoverdine production, but never by losing the trait altogether. We further show that social traits should not be studied in isolation, as they are connected through an intricate regulatory network (Balasubramanian et al. 2013). Selection for changes in one trait, such as the loss of quorum sensing, affect other traits, such as increased pyoverdine production. An integrative approach that considers the multifaceted social profiles of bacteria is needed to fully understand the evolution of sociality in these microbes.

## Acknowledgments

The authors would like to thank Carla Bello-Cabrera and Michael Baumgartner for bioinformatics assistance, and Jos Kramer for statistical advice.

