## Supplementary Figures and Tables 2-5 for "Ecology drives the evolution of diverse siderophore-production strategies in the opportunistic human pathogen *Pseudomonas aeruginosa*"

**Supplementary Material**

This file contains to following supplementary material:

- Supplementary Figures 1 - 4
- Supplementary Tables 2 - 5

Please note that the supplementary Table 1 and Tables 6 - 10 are provided in a separate spreadsheet file.

### Supplementary Figures

#### Figure S1

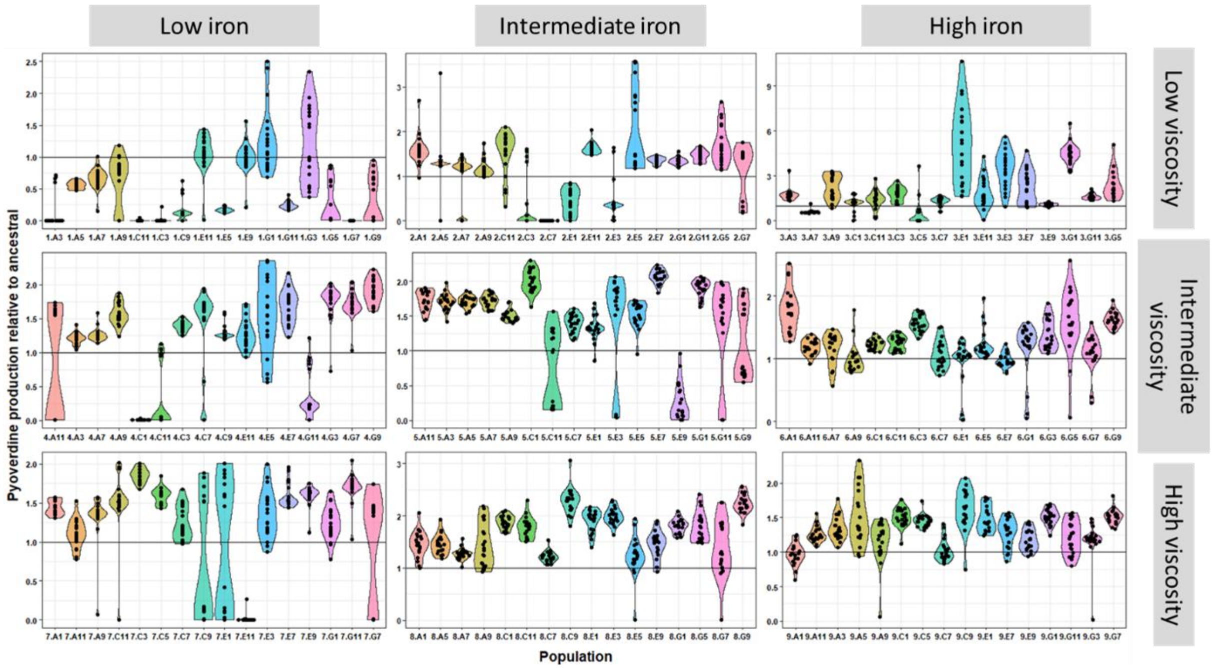


Fig. S1 – Distribution of pyoverdine production of evolved clones by experimental evolution environment, and within each environment by population. All pyoverdine production values are scaled relative to the production of the ancestral in the respective environment (represented at y=1 solid black line). Violin plot illustrate the probability of distribution of pyoverdine production phenotypes, within each population.

**Figure S2**

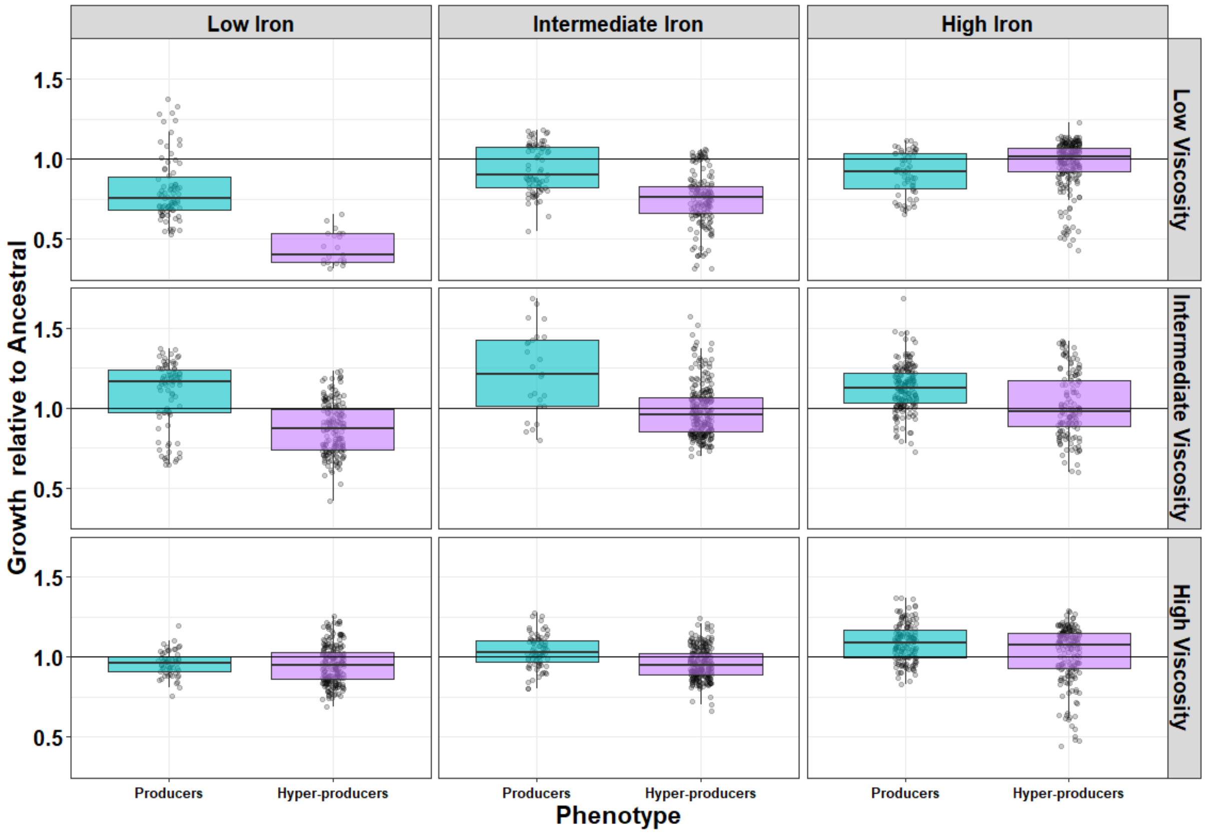


Fig S2 – Cost of being a hyper-producer. Comparison of endpoint growth ($OD_{600}$) after 48 hours between clones allocated to the producer vs hyperproducer phenotypes, separated by environments. Box plots show the median and the 1st and 3rd quartile across phenotypes, for each evolution environment (clones depicted by the individual dots). Whiskers represent the 1.5 interquartile range.

**Figure S3**

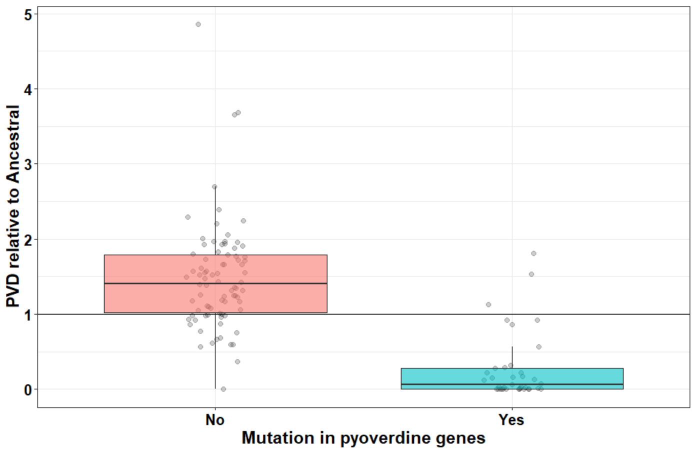


Fig. S3 – Assocation between pyoverdine production of evolved clones and the presence or absence of mutations in the pyoverdine locus. We measured pyoverdine production via its natural fluorescence, scaled to the optical density of the population (OD at 600 nm after 48 hours). Box plots show the median and the 1st and 3rd quartile across the mutational profile of each clone (depicted by the individual dots). Whiskers represent the 1.5 interquartile range. Horizontal line at y = 1 represents the ancestral phenotype.

Figure S4

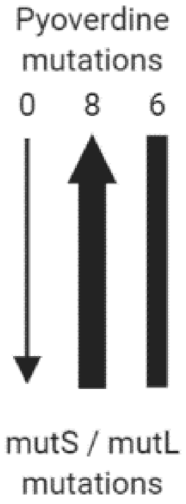


Fig. S4 – Inference of the order of mutations in the pyoverdine and mutS/mutL loci. We found that in eight populations, mutS/mutL mutations preceeded mutations in the pyoverdine locus, while there was no case in which mutS/mutL mutations arose after the spread of mutations in the pyoverdine locus. In six populations, we were unable to unambiguously establish the order of mutations.

**Supplementary Tables**

Table S2 – Summary statistics for the effects of iron availability and media viscosity on growth and pyoverdine production in the ancestral wild type.

| Response Variable | Predictor Variable | dF | F | p |
|---|---|---|---|---|
| $OD_{600}$ | Iron:Viscosity | 4:135 | 55.42 | < 0.0001 |
| | Iron | 2:135 | 228.22 | < 0.0001 |
| | Viscosity | 2:135 | 132.66 | < 0.0001 |
| Log(Pyoverdine/$OD_{600}$) | Iron:Viscosity | 4:135 | 237.67 | < 0.0001 |
| | Iron | 2:135 | 1855.82 | < 0.0001 |
| | Viscosity | 2:135 | 719.26 | < 0.0001 |

Table S3 – Summary statistics for the effects of iron availability and media viscosity on the coefficient of variation of pyoverdine production in the evolved populations.

| Response Variable | Predictor Variable | df | F | p |
|---|---|---|---|---|
| Log(Coefficient of Variation) | Iron:Agar | 4:135 | 0.2186 | 0.927 |
| | Iron | 2:139 | 4.851 | 0.009 |
| | Agar | 2:139 | 11.336 | <0.0001 |

Table S4. Summary statistics for the effect of pyoverdine production (producer vs hyper-producer) on growth of evolved clones, separated by environments.

| Treatment | dF | t | p |
|---|---|---|---|
| Low iron – low viscosity | 90.85 | -10.19 | **< 0.0001** |
| Intermediate iron – low viscosity | 229.54 | -7.807 | **< 0.0001** |
| High iron – low viscosity | 267.77 | 1.865 | 0.0632 |
| Low iron – intermediate viscosity | 248.06 | -4.689 | **< 0.0001** |
| Intermediate iron – intermediate viscosity | 264.35 | -8.412 | **< 0.0001** |
| High iron – intermediate viscosity | 306.57 | -4.37 | **0.0002** |
| Low iron – high viscosity | 264.97 | -2.815 | **0.0052** |
| Intermediate iron – high viscosity | 302.91 | -3.795 | **< 0.0001** |
| High iron – high viscosity | 313.89 | -4.386 | **0.0001** |

Table S5.

Summary statistics for how the environment in which clones evolved in, and the amount of pyoverdine produce by evolved clones, affect the extent to which pyoverdines from evolved producers stimulate the growth of ancestral non-producers.

| Response Variable | Predictor Variable | $\chi_2$ | dF | p |
|---|---|---|---|---|
| Pyoverdine/OD | Iron:Viscosity | 26.35 | 4 | < 0.0001 |
| | Iron | 7.62 | 2 | 0.0221 |
| | Viscosity | 7.02 | 2 | 0.0298 |
| | Change in pyoverdine production relative to ancestral | 278.31 | 1 | < 0.0001 |
